# Effect of mosquito saliva from distinct species on human dermal endothelial cell function *in vitro* and West Nile virus pathogenesis *in vivo*

**DOI:** 10.1101/2025.03.06.641838

**Authors:** Imke Visser, Vincent Vaes, Peter van Run, Eleanor M. Marshall, Lars Vermaat, Charlotte Linthout, Dick H.W. Dekkers, Jeroen A.A. Demmers, Marion P.G. Koopmans, Constantianus J.M. Koenraadt, Melanie Rissmann, Barry Rockx

## Abstract

During probing and feeding, an infected mosquito injects both virus and saliva into the host skin. The presence of mosquito saliva in the skin increases arbovirus pathogenesis in the bitten host, however the exact mechanism behind this remains to be determined. It is hypothesized that disease enhancement is dependent on the function of the dermal endothelium, where an increased permeability aids in the influx of virus-susceptible cells to the bite site and therefore more cells for the virus to replicate in. Here, we investigate the effects of saliva from *Culex* and *Aedes* species on the human dermal endothelial cell function *in vitro*. Furthermore, we investigate the effect of *Culex* saliva on West Nile virus (WNV) pathogenesis in a mouse model. We found that salivary gland extract from anthropophilic mosquito species (*Aedes* and *Cx. pipiens molestus*) induce permeability of the human dermal endothelium, while an ornithophilic mosquito species (*Cx. pip. pipiens*) does not. We identified that this effect is likely due to the presence of protease(s) in *Cx. pipiens molestus* saliva that are absent in *Cx. pipiens pipiens* saliva. In addition, we show that the presence of *Culex* saliva at the WNV inoculation site *in vivo* leads to more consistent weight loss, increased permeability in the inoculation site, and increased mortality compared to inoculation of WNV alone. Identification and characterization of novel salivary proteins from similar but genetically distinct mosquito species will advance the development of intervention methods to combat potential transmission risks and disease severity of emerging mosquito-borne pathogens.

## Introduction

Arthropod-borne (arbo) virus infections represent a significant global public health challenge, with approximately 3.9 billion individuals residing in tropical and sub-tropical regions at risk^1^. The transmission dynamics of arboviruses are characterized by complex ecological interactions involving a sylvatic maintenance cycle, reservoir hosts, and spill-over events mediated by bridge vector species^2^. The expanding geographical distribution of vectors, due to changing environmental and anthropogenic factors, is progressively altering arboviral transmission landscapes^3^. Notably, mosquito vector species are establishing populations in previously non-endemic regions, facilitating potential virus transmission cycles through pre-existing or novel vector-virus combinations. For example, the invasive mosquito *Aedes* (*Ae.*) *albopictus* is continuously introduced and establishing in more northern European latitudes and consequently poses a risk to public health, since this mosquito species is a competent vector for medically relevant arboviruses including dengue, chikungunya, and Zika virus^4^. Concurrent with this geographical expansion of potential vectors, certain arboviruses, traditionally confined to equatorial and Mediterranean regions, are incrementally extending their geographic distribution. These viruses are leveraging widespread vector species (spp.), such as *Culex* (*Cx.*) mosquitoes, to establish transmission cycles. West Nile virus (WNV), vectored by *Cx.* spp., exemplifies this, as it is continuously dispersing northward through European countries. Specifically, the northward expansion of WNV is evidenced by its first detection in birds and mosquitoes in the Netherlands in 2020^5^, followed by the first identification of autochthonous human infections causing neuroinvasive disease^6^.

Experimental virus infection models frequently neglect a crucial aspect of arbovirus transmission: the mosquito ‘bite’. During probing for a blood meal, mosquitoes inject saliva containing viral particles and bioactive compounds that modulate the skin’s immunological environment^7^. Animal studies demonstrate that the presence of mosquito saliva at the bite site significantly impacts viral pathogenesis, inducing an increased viral load in the skin, enhanced viremia, and higher host mortality. This phenomenon is associated with a saliva-induced increase in permeability of the dermal microvascular endothelial barrier, which allows an increased influx of immune cells, that are susceptible to arbovirus infection, to the dermis. The presence of these susceptible immune cells at the primary site of virus replication may then allow for increased viral dissemination^8^. *Ae*. *aegypti* has been the predominant mosquito species investigated for its salivary effects, primarily due to its critical role in transmitting medically important arboviruses. However, the ongoing emergence of novel arbovirus strains and transmission cycles in previously non-endemic areas necessitates understanding of how saliva from diverse mosquito species affects viral replication. While *Cx.* spp. mosquitoes— including various *Cx. pipiens* biotypes—are geographically widespread, few studies have examined how saliva from *Cx.* spp. enhances WNV pathogenesis^9,10^.

This study aimed to investigate the impact of mosquito saliva from different endemic (*Cx.* spp.) and invasive (*Ae.* spp.) species on human dermal endothelial permeability using an *in vitro* transwell system, and examine the pathogenesis-modulating potential of saliva from two distinct *Cx. pipiens* biotypes in a WNV mouse model. Characterization of mosquito salivary components and their role in arbovirus pathogenesis could facilitate the development of innovative, mosquito-focused vaccine strategies and treatments.

## Methods

### Cell culture and virus

Vero cells (African green monkey kidney epithelial cells, ATCC CCL-81) were maintained at 37°C, 5% CO_2_ in Dulbecco’s Modified Eagle’s Medium (DMEM; Lonza) supplemented with 10% heat-inactivated foetal calf serum (FCS; Sigma-Aldrich), 100U/mL penicillin, 100μg/mL streptomycin, 2mM L-glutamine (Lonza), 0.75% sodium-bicarbonate (Lonza) and 10mM HEPES buffer (Lonza). Primary Human Dermal Microvascular Endothelial cells (DMECs; Cell Systems, ACBRI 538) were maintained at 37°C, 5% CO_2_ in MV2 medium (Promocell, C-22121). Culture flasks were pre-coated with 1% porcine-derived gelatin (Sigma-Aldrich, 15-30 minutes at 37°C, 5% CO_2_). DMECs were subcultured according to the manufacturer’s protocol. For *in vivo* work, WNV-NL20 (GenBank accession number OP762595.1, EVAg 010V-04311), as described previously^11^, was used.

### Mosquito salivary gland extract

Female *Ae. aegypti* (Rockefeller strain) and *Ae. albopictus* (Agropolis strain) were laboratory-reared, whereas *Ae. japonicus*, *Cx. pipiens molestus*, and *Cx. pipiens pipiens* were field-caught and reared as described previously^12,13^. To obtain salivary gland extract (SGE), mosquitoes were dissected 7-10 days post-hatching according to a previously established protocol^14^, with few adaptations: for *in vitro* application, a concentration of one salivary gland pair (SGP) per 10μL of DMEM supplemented with 10mM HEPES buffer (Lonza), 10% FCS (Sigma-Aldrich), 50μg/mL gentamycin, 2,5μg/mL amphotericin B, 100U/mL penicillin, and 100μg/mL streptomycin was maintained. For *in vivo* application, a concentration of one SGP per 5uL of 1xPBS supplemented with 50μg/mL gentamycin, 2,5μg/mL amphotericin B, 100U/mL penicillin, and 100μg/mL streptomycin was used. To obtain SGE, the collected SGPs were sonicated using a waterbath (Distrilab) with the following settings: 4 bursts of ∼20 seconds at 220-450V with 1 minute of cooling on ice in between bursts. After sonicating, tubes were spun at 5000xg at 4°C for 10 minutes and supernatant (the SGE) was aliquoted. In all experiments, an equivalent of one SGP (containing ∼1.8μg protein) per replicate or animal was used unless stated otherwise.

### Trans-Endothelial Electrical Resistance assay

DMECs (passage 8-10) were seeded into the apical compartment of 0.4μm pore 12mm diameter 12-well transwell inserts (Corning, 3460), pre-coated with 1% porcine gelatin, at a density of 1.8×10^5^ cells per well in 800μL MV2 medium. Transwells were incubated at 37°C, 5% CO_2_. At 2 days post-seeding, MV2 medium on both the apical and basolateral side was refreshed. At 4 days post-seeding (t0), prior to adding SGE, the barrier function was assessed by measuring the trans-endothelial electrical resistance (TEER) using the EVOM2 epithelial voltohmmeter and STX2 handheld chopstick electrodes (World Precision Instruments) placed in the basolateral and apical compartment of the transwells. After t0 measurement, SGE of each mosquito species was added to the medium in the apical compartment of the transwell. As a positive control for increased endothelial permeability, 100ng/mL of recombinant human TNFα (Promocell) was added to untreated DMECs. TEER measurements were performed every 3 hours post-saliva stimulation (hps) up until 12 hps. At 24 hps, TEER was measured and then medium from the apical and basolateral compartment was refreshed. At 48 hps, TEER was measured again to assess recovery of the endothelial barrier to baseline. TEER values were normalised to a gelatin coating-only control and surface area, obtaining Ohm/cm^2^ values. The relative TEER was further calculated as the percentage resistance of SGE-treated transwells divided by the untreated control transwells, obtaining %TEER values. To assess the involvement of salivary proteins in endothelial barrier permeability, SGE was heat-inactivated at 65°C for 30 mins prior to adding SGE to the DMECs. To assess the involvement of salivary proteases in endothelial barrier permeability, protease inhibitor (cOmplete Mini EDTA-free Protease Inhibitor Cocktail, Roche) was added to the culture medium prior to adding SGE.

### Mass-spectrometry of *Culex* saliva

For global proteome analysis, 40 SGPs (containing 75μg total protein as determined by BCA protein assay) per *Cx. pipiens* biotype were collected in PBS containing cOmplete Mini EDTA-free Protease Inhibitor Cocktail (Roche). Saliva was collected from the SGPs by sonication in a water bath combined with 3 freeze-thaw cycles followed by centrifugation to harvest saliva proteins in the supernatant. Isolated proteins were reduced in lysis buffer with 5mM dithiothreitol and alkylated with 10mM iodoacetamide. Next, proteins were digested with 2.5μg trypsin (1:40 enzyme:substrate ratio) overnight at 37°C. After digestion, peptides were acidified with trifluoroacetic acid (TFA) to a final concentration of 0.5% and centrifuged at 10,000g for 10 min to spin down the precipitated SDC. Peptides in the supernatant were desalted on a 50mg C18 Sep-Pak Vac cartridge (Waters). After washing the cartridge with 0.1% TFA, peptides were eluted with 50% acetonitrile and dried in a Speedvac centrifuge. Peptides were then analyzed by nanoflow LC-MS/MS as described previously^15^. For data analysis, DIA raw data files were analyzed with the Spectronaut Pulsar X software package (Biognosys, version 18.0.230605.50606) using directDIA for DIA analysis including the IDPicker algorithm for protein inference, using a combined protein reference fasta database containing all *Cx. pipiens* protein sequences (taxonomy ID 7175; containing 75,763 entries). The mass spectrometry proteomics data have been deposited to the ProteomeXchange Consortium via the PRIDE^16^ partner repository with the dataset identifier PXD060865.

### Ethics

All animal procedures were performed in compliance with the Dutch legislation for the protection of animals used for scientific purposes (2014, implementing EU Directive 2010/63) and other relevant regulations. The licensed establishment where this research was conducted (Erasmus MC) has an approved OLAW Assurance #A5051-01. Animal experiments were conducted under a project license from the Dutch competent authority and the study protocol #2010606 was approved by the institutional Animal Welfare Body.

### Intradermal inoculation of WT mice in the presence and absence of *Culex* saliva

To assess the effect of mosquito saliva on WNV pathogenesis, specific pathogen-free wild-type (WT) C57BL/6 age-matched female mice were obtained from Charles River Laboratories and inoculated intradermally with 10^5^ TCID_50_/mouse WNV in the presence or absence of *Cx. pipiens molestus* (MOL) or *Cx. pipiens pipiens* (PIP) SGE. The experiment was conducted in two independent rounds: round one comprised MOL, WNV, and WNV+MOL groups, while round two included PIP, WNV, and WNV+PIP groups, i.e. each with its own WNV control group. Animals were injected into the footpad of the right hindleg with a total volume of 15μL as follows; WNV-only groups: 10μL WNV stock + 5μL PBS supplemented with antibiotics and antifungals (see previous section ‘mosquito salivary gland extract’); SGE-only groups: 10μL PBS + 5μL SGE; WNV+SGE groups: 10μL WNV stock + 5μL SGE. Groups of animals were euthanized at 4 hours post-inoculation (hpi) and 24 hpi to investigate the (early) effects of SGE on vascular permeability. Prior to euthanasia, 100μL (2.5mg/mouse) of 70kDa Fluorescein isothiocyanate-dextran (FITC-dextran) (Sigma-Aldrich) was injected intravenously into the tail vein and allowed to circulate for 1 hour. The remaining animals were monitored for clinical signs of disease and weighed daily. To study viremia kinetics, a 30μL blood sample was collected from the tail vein every other day starting at 2 dpi up until 10 dpi. At the end of the experiment (10 dpi), or when reaching humane end-points, animals were euthanized and various visceral and brain tissues were harvested. Sample processing, serum RNA isolation, qPCR, and virus (tissue) titrations were performed as previously described^11^.

### FITC-dextran histochemistry staining and quantification of mouse footpads

During necropsy, the feet from both the right and the left hindleg were collected and fixed in 4% buffered formaldehyde. After fixation, the feet were incubated for a minimum of 14 days in 10% EDTA to allow decalcification, and were subsequently processed and embedded in paraffin and cut into 3-5μm thick sections. To visualize the FITC-dextran in the feet, slides were deparaffinized and incubated with polyclonal Rabbit anti-FITC/HRP (DAKO, Agilent). Slides were subsequently incubated in Biotin Amplification Reagent following manufacturer’s protocol (TSA PLUS Biotin KIT, PerkinElmer) followed by a streptavidin-HRP conjugate (DAKO, Agilent).

FITC-dextran-stained foot slides were scanned using the Nanozoomer 2.0 HT (Hamamatsu) and scans were imported to QuPath-0.3.2 to obtain representative pictures per footpad for further quantification of FITC signal. As the effect of saliva is local^9^, data obtained from the inoculated right hindfoot was then normalised to the signal present in the non-inoculated left hindfoot of the same animal to account for inter-animal differences in vessel permeability. FITC signal was most consistent in the muscle areas of the footpads across all groups, therefore three representative pictures of the muscle area of each footpad were made, avoiding major vessel areas, to quantify FITC signal. FITC signal was quantified using the pixel classification thresholding function of QuPath version 0.3.1^17^. A threshold of 0.15 was used for all images.

### Statistical analysis

All statistical analyses, indicated in figure legends, were performed using Graphpad Prism version 10.4.1.

## Results

### SGE from distinct *Aedes* species and *Culex pipiens molestus* induce permeability of human dermal endothelium *in vitro*

To investigate mosquito species-specific effects of mosquito saliva on human dermal endothelium permeability, we measured the TEER in DMECs following stimulation with SGE. SGE from *Ae. aegypti, Ae. albopictus*, and *Ae. japonicus* significantly decreased TEER values at 6-12 hps, indicating increased endothelial permeability (**Fig.1A**). While *Cx. pip. molestus* SGE induced significant permeability between 3-12 hps, *Cx. pip. pipiens* SGE showed no effect on TEER values at any timepoint (**Fig.1B**). Notably, protein concentrations of the two SGE preparations was comparable and increasing SGE concentration did not alter these responses for either *Cx. pip. molestus* or *Cx. pip. pipiens* (**Fig.S1**), suggesting that the lack of effect on permeability of *Cx. pip. pipiens* SGE is not due to a differences in total protein concentration.

**Figure 1.**
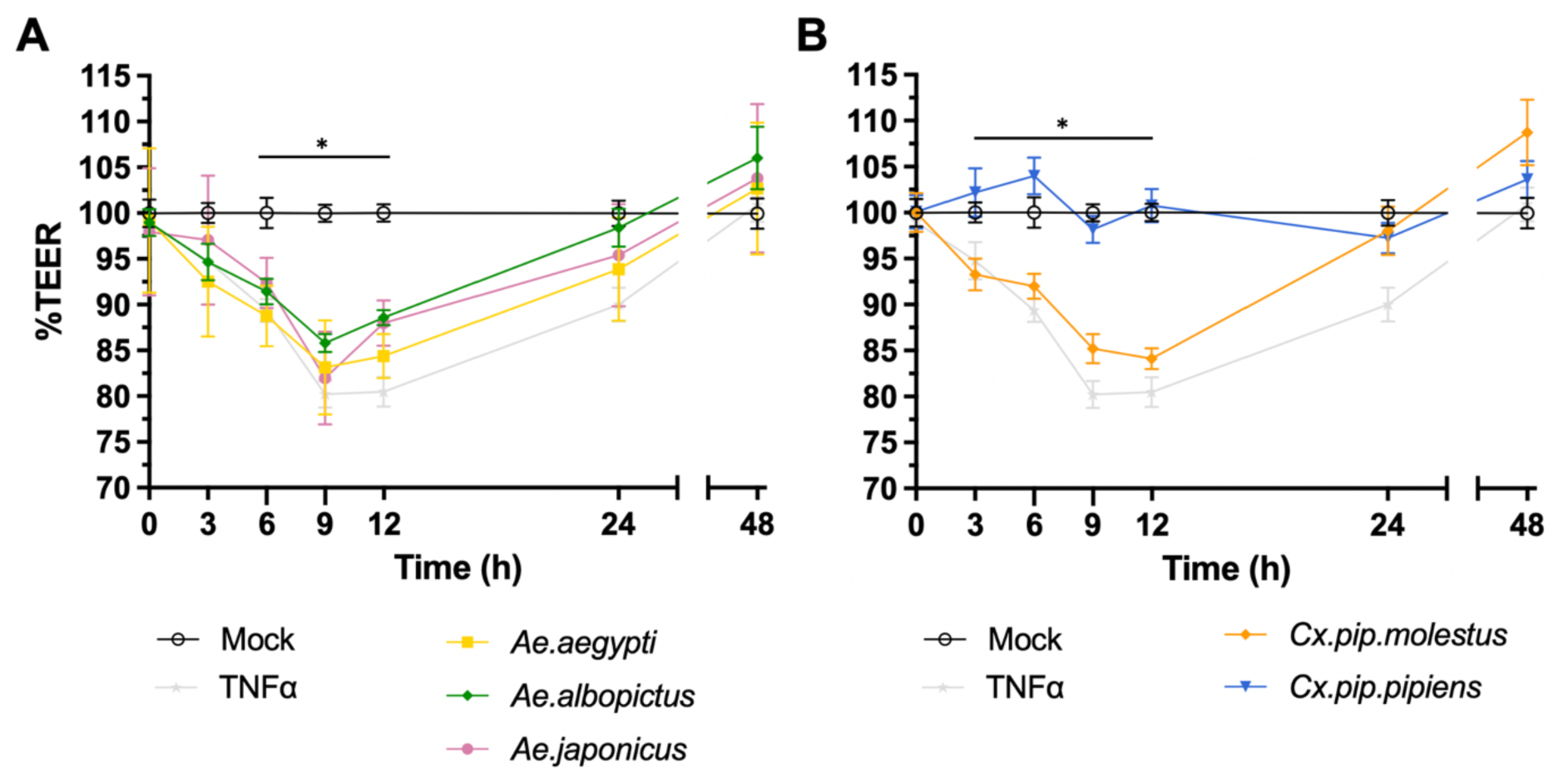
Effect of SGE on the barrier function of human dermal endothelium. Error bars represent the SEM. Trans-endothelial electrical resistance (TEER) of primary human dermal microvascular endothelial cells (DMECs) grown on the apical side of a transwell insert was measured after stimulation of DMECs with SGE (1 SGP per well) or TNF-α (100ng/mL) as a positive control. Data pooled from three individual experiments, three transwells per condition, each measured three times per timepoint. Statistically significant differences between SGE and mock are indicated with an asterisk \**p*<0.05; as measured by 2way ANOVA multiple comparisons **(A)** DMECs were stimulated with SGE from either *Ae. aegypti*, *Ae. albopictus*, *Ae. japonicus*, or **(B)** *Cx. pipiens molestus* or *Cx. pipiens pipiens*. The %TEER was measured every 3 hours over a time-course of 12 hours, and again at 24 and 48 hours post-stimulation.

### Proteases in *Culex pipiens molestus* SGE are associated with increased endothelial permeability

Given previous reports of *Ae. aegypti* salivary peptides mediating endothelial permeability^8^, we examined the potential role of proteins in *Cx. pip. molestus* (MOL) SGE through heat-inactivation. Heat-treatment of MOL SGE completely abrogated the induction of endothelial permeability at 9 hps (**Fig.2A**), suggesting that a protein is involved in this process. Therefore, we subsequently performed a pilot mass spectrometry experiment of SGE from both *Cx. pipiens* biotypes to potentially identify proteins that are uniquely present in MOL SGE. This initial screen revealed the presence of several serine proteases in MOL but not *Cx. pip. pipiens* (PIP) SGE (**Table.S1**). To confirm this finding, we show that the addition of protease-inhibitors to MOL SGE abolished its permeability-inducing effect on DMECs (**Fig.2B**). Overall, we observed that SGE from MOL induced permeability to the same extent as SGE from the three *Aedes* species, whereas PIP SGE did not induce permeability, and that this could be due to the presence of proteases in MOL SGE that are absent in PIP SGE.

**Figure 2.**
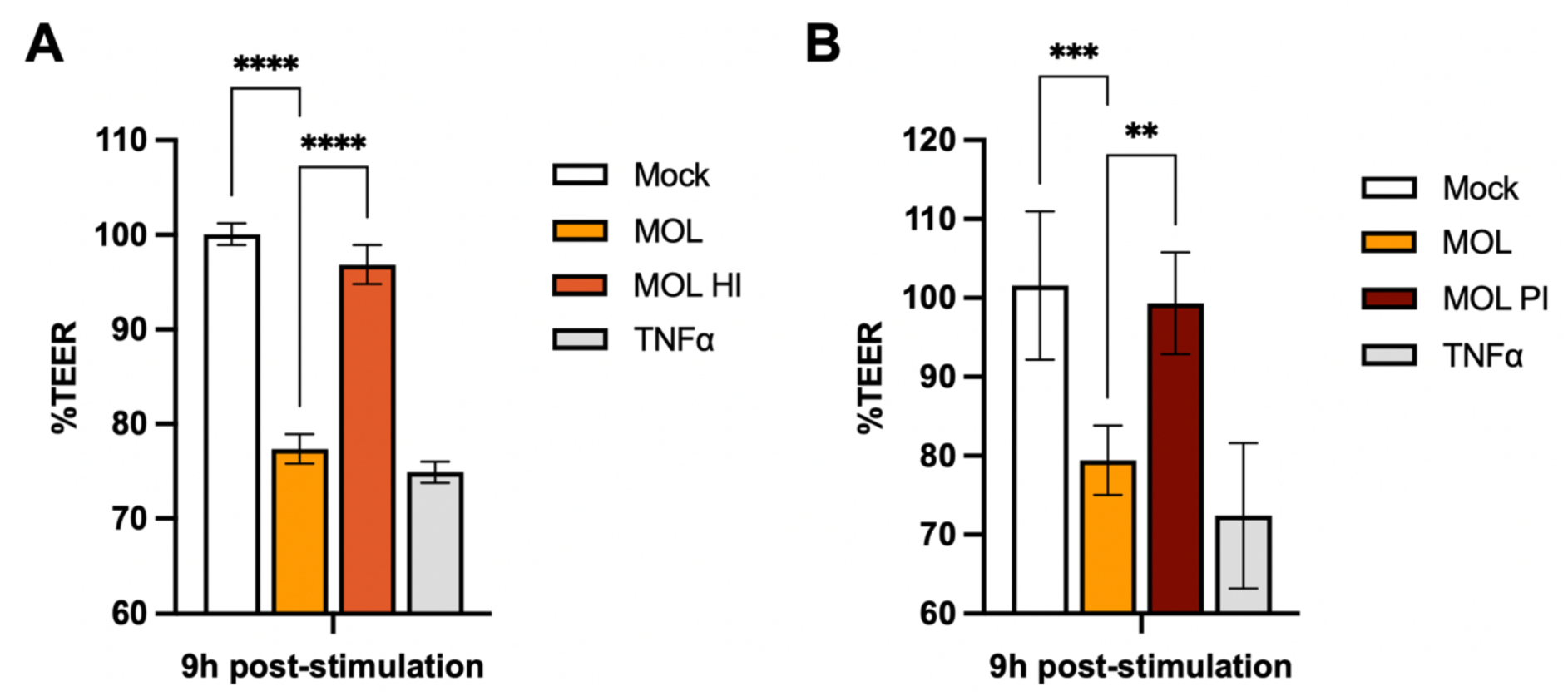
Effect of heat-inactivation and protease-inhibition of *Cx. pip. molestus* SGE. Error bars represent the SEM. **(A)** SGE from *Cx. pipiens molestus* (MOL) was heat-inactivated (MOL HI) and %TEER was measured at 9 hours post-stimulation. Data pooled from three individual experiments, three transwells per condition, each measured three times per timepoint. **(B)** MOL SGE was treated with a protease-inhibitor (MOL PI) and %TEER was measured at 9 hours post-stimulation. Data pooled from two individual experiments, three transwells per condition, each measured three times per timepoint. Statistically significant differences are indicated by asterisks as follows: \*\**p*<0.005; \*\*\**p*<0.001; \*\*\*\**p*<0.0001 as measured by Ordinary one-way ANOVA multiple comparisons.

### SGE from *Culex pipiens* biotypes enhance WNV-induced morbidity and mortality in mice

An increased permeability at the mosquito bite site has been previously speculated to be the main driver of enhanced arbovirus pathogenesis^8^. Based on our *in vitro* data, we therefore hypothesized that SGE from MOL and PIP would differentially affect arbovirus pathogenesis *in vivo*. To investigate the influence of saliva from different *Cx. pipiens* biotypes on WNV pathogenesis, we inoculated C57BL/6 wild-type mice with WNV either alone or in the presence of MOL or PIP SGE. In order to assess whether the addition of saliva would affect mortality or alter any WNV-related clinical symptoms, an inoculation dose of 10^5^ TCID_50_/mouse was chosen as this dose was previously shown to result in 40% probability of survival in mice^11^. Disease progression was monitored over 10 days (**Fig.3A**) with SGE-only inoculations serving as negative controls.

**Figure 3.**
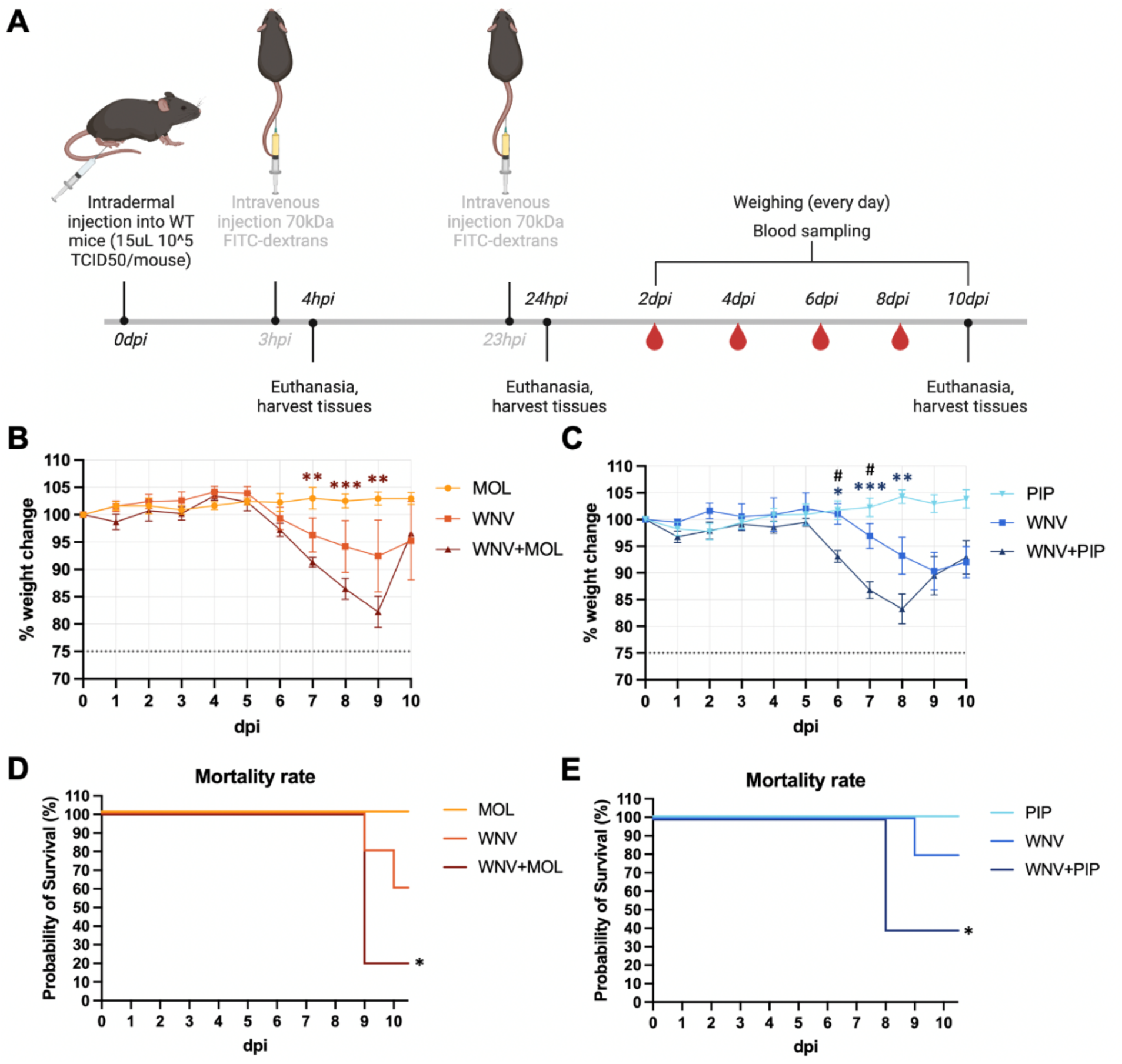
Effect of *Cx. pipiens molestus* (MOL) and *Cx. pipiens pipiens* (PIP) SGE on weight loss and mortality in mice post-WNV inoculation. Error bars represent the SEM. **(A)** Schematic overview of the timeline of the experiment. **(B)** Weight loss kinetics of animals inoculated with either MOL, WNV, or a combination of WNV+MOL over a time-course of 10 days post-inoculation (dpi). Statistically significant differences between MOL and WNV+MOL are indicated by asterisks \*\**p*<0.005 and \*\*\**p*<0.001 as tested by 2way ANOVA Tukey’s multiple comparisons. **(C)** Weight loss kinetics of animals inoculated with either PIP, WNV, or a combination of WNV+PIP over a time-course of 10 dpi. Statistically significant differences between PIP and WNV+PIP are indicated by asterisks \**p*<0.05; \*\**p*<0.005; \*\*\**p*<0.001 and between WNV and WNV+PIP ^#^*p*<0.05 as tested by 2way ANOVA Tukey’s multiple comparisons. **(D)** Survival curve of animals inoculated with MOL, WNV, or WNV+MOL. **(E)** Survival curve of animals inoculated with PIP, WNV, or WNV+PIP. Significant differences in survival between MOL and WNV+MOL, and PIP and WNV+PIP are indicated by asterisk \**p*<0.05 tested by Log-rank Mantel-Cox test.

Mice inoculated with WNV or WNV+MOL began losing weight at 6 days post-inoculation (dpi), continuing through 9 dpi when several animals reached humane endpoints. While the WNV group showed no significant weight loss compared to the MOL group, the WNV+MOL group exhibited significant weight loss between 7-9 dpi compared to MOL controls (**Fig.3B**). Similarly, in the PIP experiment, both WNV and WNV+PIP groups showed weight loss beginning at 6 dpi. However, only the WNV+PIP group demonstrated significant weight loss between 6-8 dpi compared to PIP controls. Moreover, the WNV+PIP group showed significantly greater weight loss at 6 and 7 dpi compared to the WNV group (**Fig.3C**), suggesting that PIP SGE also enhances WNV-induced weight loss. Survival analysis revealed significant differences between groups, where the WNV+MOL group showed markedly reduced survival (20%) compared to the MOL controls, while the survival rate of the WNV group (60%) did not significantly differ from MOL (**Fig.3D**). The same pattern was observed in the PIP experiment, where the WNV+PIP group exhibited significantly reduced survival (40%), while the WNV group showed no significantly different probability of survival (80%) compared to PIP controls (**Fig.3E**). Overall, these data show that addition of MOL or PIP SGE to the WNV inoculum enhances WNV-induced weight loss and mortality post-intradermal inoculation.

### *Culex pipiens molestus* SGE enhances WNV viremia in mice

To investigate potential effects on host-vector transmission dynamics, we examined whether MOL or PIP SGE influenced WNV viremia kinetics. In both WNV and WNV+MOL group, viremia peaked at 2 dpi with titers reaching 2.0-2.5 Log_10_TCID_50_eq/mL, followed by a decline at 4 dpi and a slight increase at 6 dpi, consistent with our previous WNV viremia patterns^11^. Overall viremia was significantly higher in the WNV+MOL group compared to WNV (**Fig.4A**). The PIP experiment showed similar kinetics, albeit with marginally lower peak titers (1.5-2.0 Log_10_TCID_50_eq/mL) in both WNV and WNV+PIP groups. No significant differences were detected between WNV and WNV+PIP (**Fig.4B**). These data suggest that the addition of MOL, but not PIP, saliva to the WNV inoculum increases viremia.

**Figure 4.**
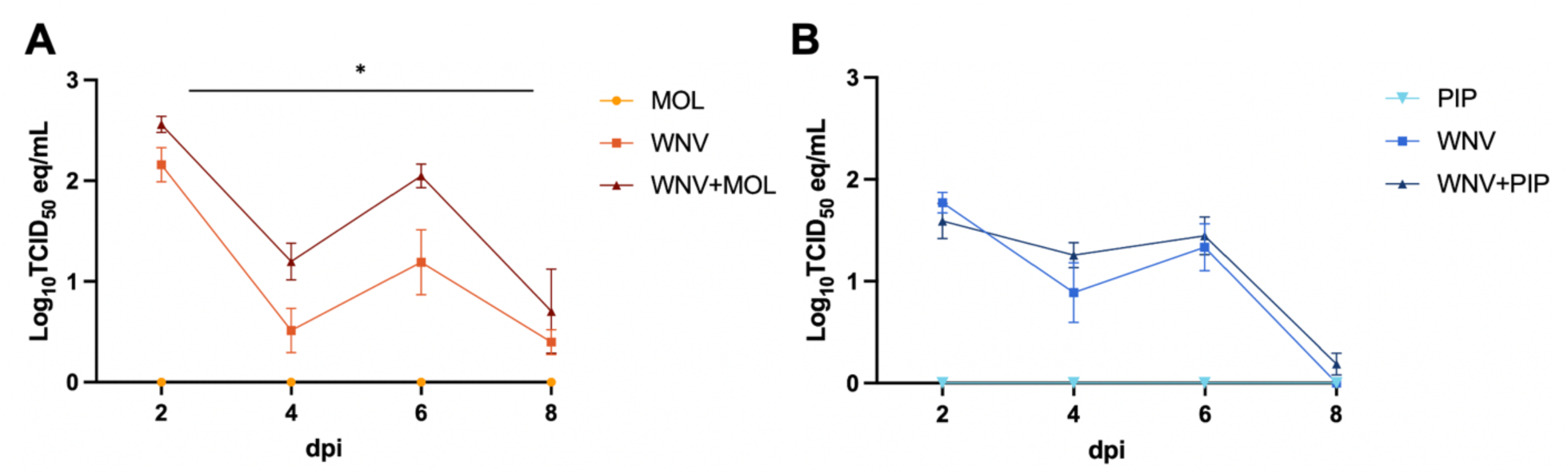
Viremia kinetics post-WNV inoculation in the presence or absence of *Cx. pipiens molestus* (MOL) or *Cx. pipiens pipiens* (PIP) SGE. Blood was taken every 2 dpi on up until 8 dpi. Error bars represent the SEM. **(A)** Viral genomic load in the blood of mice inoculated with either MOL, WNV, or WNV+MOL. Statistically significant differences between WNV and WNV+MOL are indicated by the asterisk and measured with an unpaired t-test of area under the curve (AUC) analysis. \**p*<0.05 **(B)** Viral genomic load in the blood of mice inoculated with either PIP, WNV, or WNV+PIP. No statistically significant differences were observed between WNV and WNV+PIP, as measured by unpaired t-test of AUC.

### *Culex pipiens* biotype SGE induces increased dermal vascular permeability in mice

Mosquito saliva at the inoculation site may enhance arbovirus pathogenesis by increasing dermal endothelial permeability^8^. To investigate this, and any potential biotype-specific differences as observed in our *in vitro* TEER experiment, we administered FITC-dextran intravenously and quantified its extravasation in the footpad. We compared saliva-treated and WNV-only groups at 4 hpi, i.e. before viral replication would have occurred. At 4 hpi, we observed that inoculation of MOL SGE led to a significantly increased FITC signal compared to the WNV group (**Fig.5A-B,D**). We also observed FITC signal in the WNV+MOL group (**Fig.5C**), and although the signal was slightly higher than in the WNV-only group, this difference was not statistically significant (**Fig.5D**). A similar trend was observed at 24 hpi, but the differences in FITC signal between groups were less pronounced (**Fig.5E-G**), and not significantly different (**Fig.5H**).

**Figure 5.**
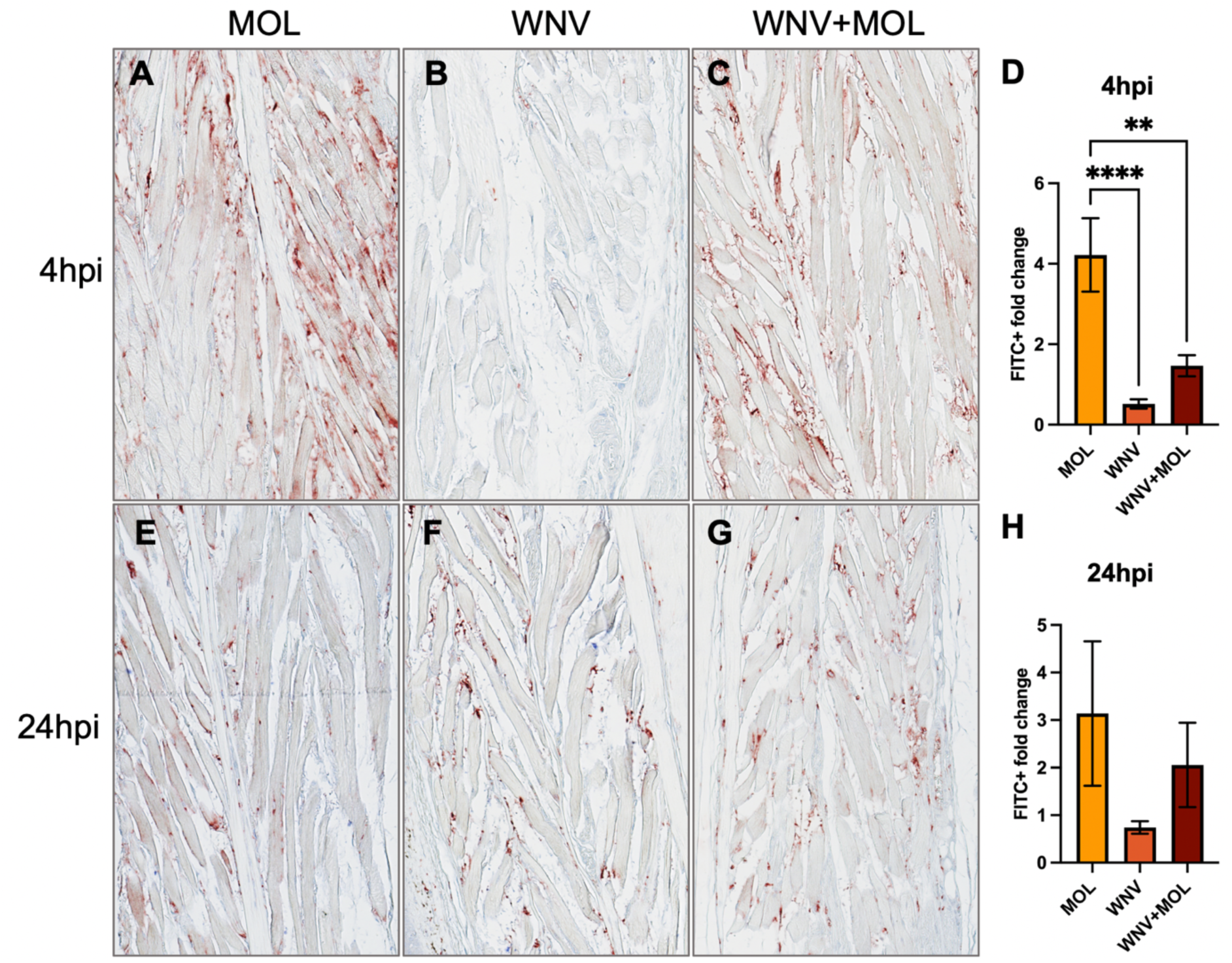
Effect of *Cx. pipiens molestus* (MOL) saliva and WNV inoculation on FITC-dextran influx into the mouse footpad at 4 and 24 hours post-inoculation (hpi). (A-C, E-G) Representative pictures (20x magnification) of FITC signal as measured by histochemistry staining in footpads of the right hindleg (inoculation site). **(D+H)** Quantification of FITC+ signal per 20x zoom footpad section. Per footpad, 3 random pictures were taken of the muscle area to represent each mouse. Signal of the right footpad (inoculation site) was divided over the signal of the left footpad (distal from inoculation site, internal negative control) to measure the FITC+ fold change. One-way ANOVA was used for statistical analysis. \*\**p*<0.005; \*\*\*\**p*<0.0001. **(A-C)** FITC signal in the footpad at 4 hpi with either **(A)** MOL, **(B)** WNV only, **(C)** WNV+MOL. **(D)** Quantification of FITC signal in the mouse footpads at 4 hpi. **(E-G)** FITC signal in the footpad at 24 hpi with either **(E)** MOL, **(F)** WNV only, **(G)** WNV+MOL.

The PIP experiment yielded similar results, with the PIP group exhibiting significantly higher FITC signal in the footpad at 4 hpi compared to the WNV group (**Fig.6A-B,D**). Although the FITC signal in the WNV+PIP group appeared to be more pronounced (**Fig.6C**), it was not significantly different from the WNV group (**Fig.6D**). At 24 hpi, the FITC signal in the PIP group remained significantly elevated relative to the WNV group (**Fig.6E-F,H**), while the FITC signal in the WNV+PIP group had diminished slightly compared to 4 hpi (**Fig.6G**). No significant differences were observed between the WNV and WNV+PIP groups at either time point (**Fig.6H**). Overall, the observed trends suggest that the addition of either MOL or PIP SGE to WNV inoculation enhances vascular permeability at both early time-points post-inoculation, relative to WNV alone.

**Figure 6.**
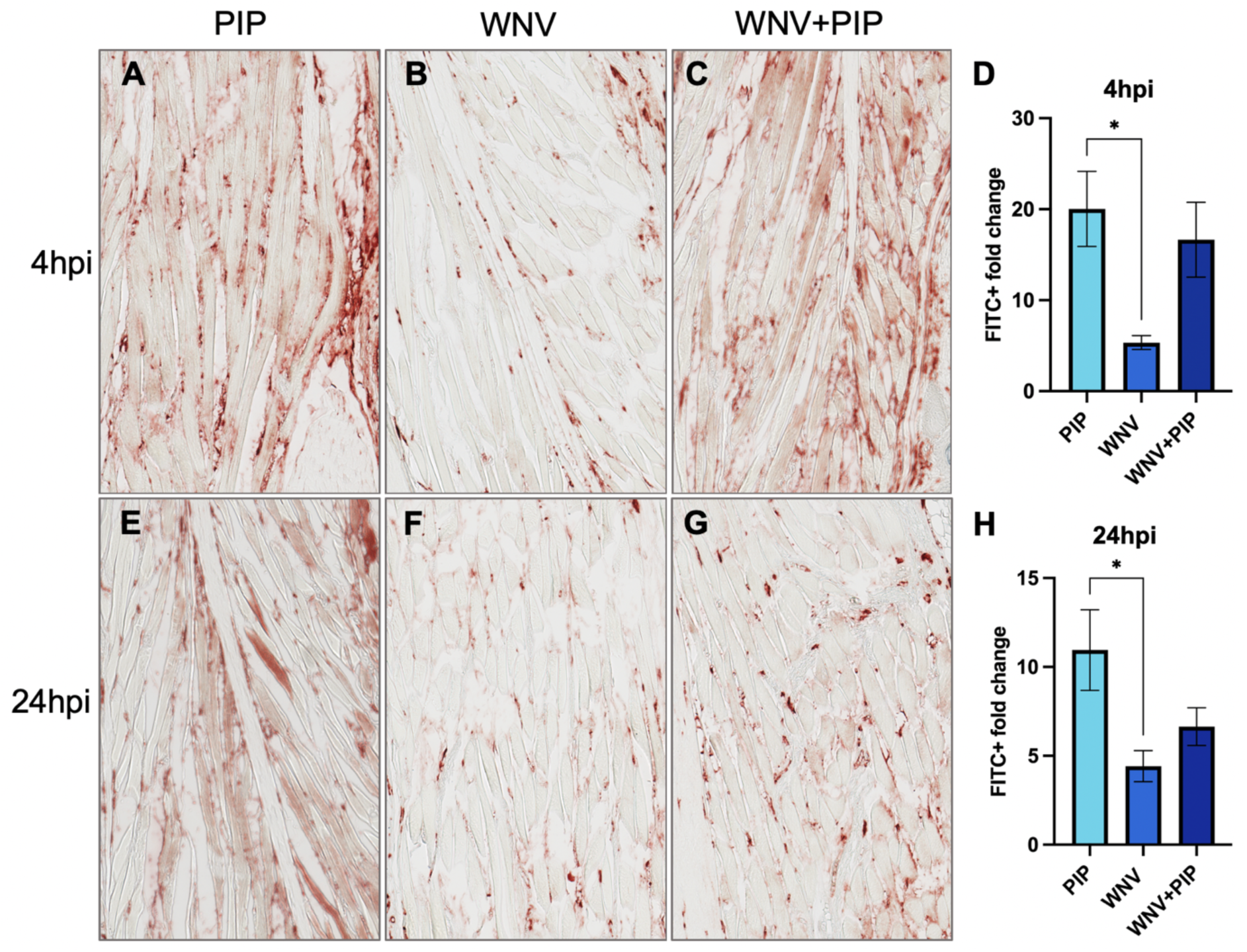
Effect of *Cx. pipiens pipiens* (PIP) saliva and WNV inoculation on FITC-dextran influx into the mouse footpad at 4 and 24 hours post-inoculation (hpi). (A-C, E-G) Representative pictures (20x magnification) of FITC signal as measured by histochemistry staining in footpads of the right hindleg (inoculation site). **(D+H)** Quantification of FITC+ signal per 20x zoom footpad section. Per footpad, 3 random pictures were taken of the muscle area to represent each mouse. Signal of the right footpad (inoculation site) was divided over the signal of the left footpad (distal from inoculation site, internal negative control) to measure the FITC+ fold change. One-way ANOVA was used for statistical analysis. \**p*<0.05. **(A-C)** FITC signal in the footpad at 4 hpi with either **(A)** PIP, **(B)** WNV only, **(C)** WNV+PIP. **(D)** Quantification of FITC signal in the mouse footpads at 4 hpi. **(E-G)** FITC signal in the footpad at 24 hpi with either **(E)** PIP, **(F)** WNV only, **(G)** WNV+PIP.

## Discussion

The recent emergence of WNV in Europe, and its first autochthonous human infections in the Netherlands in 2020^6^, underscores the urgent need to understand factors influencing WNV pathogenesis in humans. This urgency is heightened by the expanding geographical range of arbovirus vectors and the predicted intensification of arbovirus transmission dynamics, driven by complex ecological, socioeconomic, and climatic variables. Given the critical role of mosquito saliva in modulating arboviral disease outcomes in vertebrate hosts, we investigated the species-specific effects of mosquito saliva on arbovirus pathogenesis. Our study examined whether salivary gland extract (SGE) from distinct *Ae.* spp. and two *Cx. pipiens* biotypes differentially affect endothelial permeability *in vitro*. Additionally, we evaluated the impact of SGE from the two *Cx. pipiens* biotypes on WNV pathogenesis in a mouse model.

Our *in vitro* studies demonstrated that SGE from *Ae. aegypti*, *Ae. albopictus*, and *Ae. japonicus* induced human dermal endothelial permeability, consistent with previous findings using identical assays to examine *Ae. aegypti* saliva^8,18^. The comparable effects on permeability observed across these *Aedes* species likely reflect their conserved salivary proteins^19^, suggesting a shared underlying mechanism. Notably, while MOL SGE induced endothelial permeability comparable to the three *Aedes* species, PIP SGE showed no effect. These biotypes, though morphologically indistinguishable, exhibit distinct host preferences: MOL primarily feeds on mammals, whereas PIP preferentially feeds on birds^20,21^. This behavioural difference aligns with our previous hypothesis that ornithophilic mosquitoes may lack specific anti-coagulation factors present in anthropophilic species’ saliva, reflecting the slower blood-clotting mechanism in birds compared to humans^7^. We performed an initial mass spectrometry screen to compare the proteome of the two *Cx. pipiens* biotypes, and found several serine proteases present in MOL SGE that were absent in PIP SGE. The functional significance of these proteases was demonstrated when protease inhibitors abrogated the permeability-inducing effect of MOL SGE on human dermal endothelium *in vitro*. Proteases may influence host responses through multiple mechanisms. They could modulate the coagulation cascade and activate protease-activated receptors (PARs) on endothelial cells, triggering immune cell aggregation^22^. This aligns with previous observations of immune cell recruitment to the bite site as a response to mosquito saliva^8^, although a different mosquito species and salivary derivative was studied there. Furthermore, serine proteases released by activated neutrophils are shown to disrupt endothelial cytoskeletal architecture and increase permeability, an effect that is prevented by serine protease inhibition^23^. Salivary serine proteases from MOL may exert a similar effect on endothelial cell function, though further investigation is needed to elucidate their specific role in cytoskeletal disruption. Interestingly, one study found that serine proteases in *Ae. aegypti* saliva enhance Dengue virus infection *in vitro* by degrading extracellular matrix proteins. Inhibiting serine protease activity in both *Ae. aegypti* and *Cx. tarsalis* saliva significantly reduced Dengue virus and WNV infection in mouse footpads, respectively^24^.

The differential effects of MOL and PIP SGE on endothelial permeability *in vitro*, coupled with the previously reported role of serine proteases^24^ and endothelial permeability^10^ in saliva-mediated enhancement of arbovirus pathogenesis, led us to investigate the *in vivo* effects of MOL and PIP SGE. We hypothesized that MOL SGE would enhance WNV pathogenesis, while PIP SGE would not. Contrary to our hypothesis, both MOL and PIP SGE enhanced weight loss and mortality. Previous studies have demonstrated that mosquito saliva at the bite site enhances host viremia during WNV infection^9,10,25^, although the exact mechanism behind this is unknown. While we observed significant elevation in viremia in the WNV+MOL group compared to WNV alone, the WNV+PIP group showed no enhancement. Both MOL and PIP SGE significantly increased footpad permeability at 4 hpi, consistent with previous findings^18^. The greater permeability induced by MOL compared to WNV+MOL was unexpected, suggesting possible virus-saliva interactions that modulate the effects on dermal endothelial permeability. Alternatively, host defence mechanisms triggered by viral infection might counteract the permeability-enhancing effects of mosquito saliva.

While WNV+MOL and WNV+PIP groups showed a clear trend towards increased permeability at 4 hpi compared to WNV alone, this was not statistically significant. Overall, WNV+MOL and WNV+PIP inoculation resulted in more homogeneous disease outcomes in mice than WNV alone. Both WNV+MOL and WNV+PIP induced significant morbidity and mortality compared to SGE-only controls, while WNV alone resulted in more variable responses not statistically significant from SGE-only controls. This heterogeneity in WNV groups aligns with our previous observations using an identical inoculation dose and route^11^. Although differences between WNV and WNV+SGE groups were largely non-significant (except for the viremia and brief weight loss differences in the MOL and PIP experiment, respectively), the addition of SGE appeared to enhance the experimental model’s sensitivity in detecting the effects of WNV infection via intradermal inoculation. However, several methodological considerations warrant attention. Our use of SGE, rather than saliva or a bite from live infected mosquitoes, likely introduced salivary gland compounds not typically present during natural mosquito probing and feeding. Additionally, mosquito salivary gene expression varies between blood-fed versus sugar-fed, and infected versus non-infected mosquitoes^26^. Therefore, developing a more physiologically relevant model would require using SGE or live mosquito bites comparing blood-fed with sugar-fed, and infected with uninfected mosquitoes.

The discrepancy between our *in vitro* and *in vivo* findings may reflect fundamental differences between the two experimental systems. While PIP SGE had no effect on human endothelial cell permeability *in vitro*, it increased permeability in our mouse model, suggesting that *in vitro* analysis of endothelial permeability is not always predictive of *in vivo* results. This is supported by the fact that following an increased dermal permeability^8^, the influx of immune cells to the bite site is an important factor in disease enhancement *in vivo*^27^, as mentioned previously, which is not modelled in our *in vitro* endothelial barrier assay. Moreover, our *in vitro* model consists of a DMEC monolayer which does not account for any effects on the immune responses of skin-resident cells upon infection or SGE-stimulation, that could cause an indirect activation of the endothelium. For example, infected keratinocytes release a range of chemokines and cytokines involved with immune cell recruitment towards the site of infection^7^. Establishment of a co-culture system consisting of DMECs and, for example, dermal fibroblast, epidermal keratinocytes, and Langerhans cells to study direct and indirect effects of both SGE and virus infection could help elucidate the differences observed between *in vitro* and *in vivo* experiments. The differential *in vitro* versus *in vivo* results may also stem from organism-specific differences, and could be investigated further by a comparative analysis of PIP SGE effects on human versus mouse dermal endothelial cells using our *in vitro* assay.

In conclusion, both *Ae.* spp. and MOL SGE induced permeability in human dermal endothelium *in vitro*, whereas PIP SGE showed no effect. This effect is likely mediated by proteases in MOL SGE. Our *in vivo* experiments revealed that co-inoculation of WNV with either MOL or PIP SGE enhanced disease outcomes, including increased morbidity, mortality, and permeability at the inoculation site, compared to inoculation with WNV alone.

## Supplementary

**Supplementary Figure 1.**
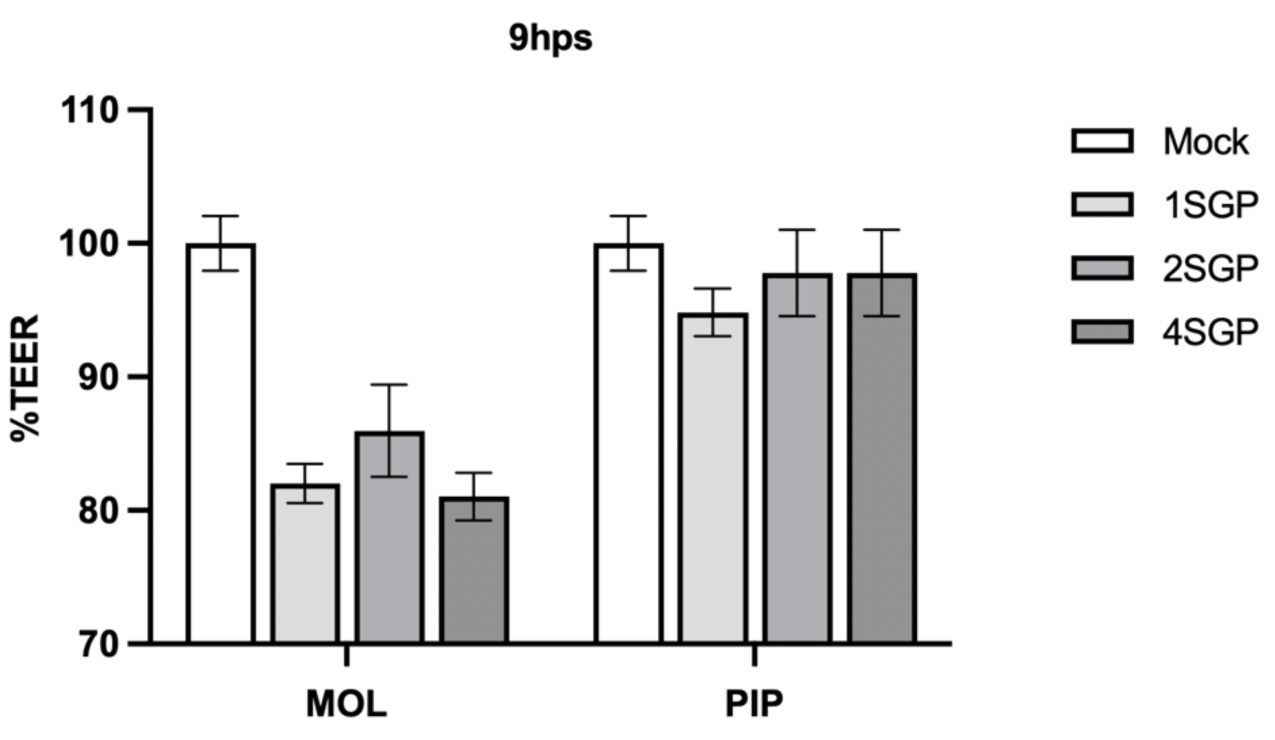
Effect of increased salivary gland pair (SGP) concentration on the trans-endothelial electrical resistance (TEER). An equivalent of either 1, 2, or 4 *Cx. pipiens molestus* (MOL) or *Cx. pipiens pipiens* (PIP) SGP was added to primary human dermal microvascular endothelial cells after which TEER was measured at 9 hours post-stimulation (hps). Data from one individual experiment.

**Supplementary Table 1.**
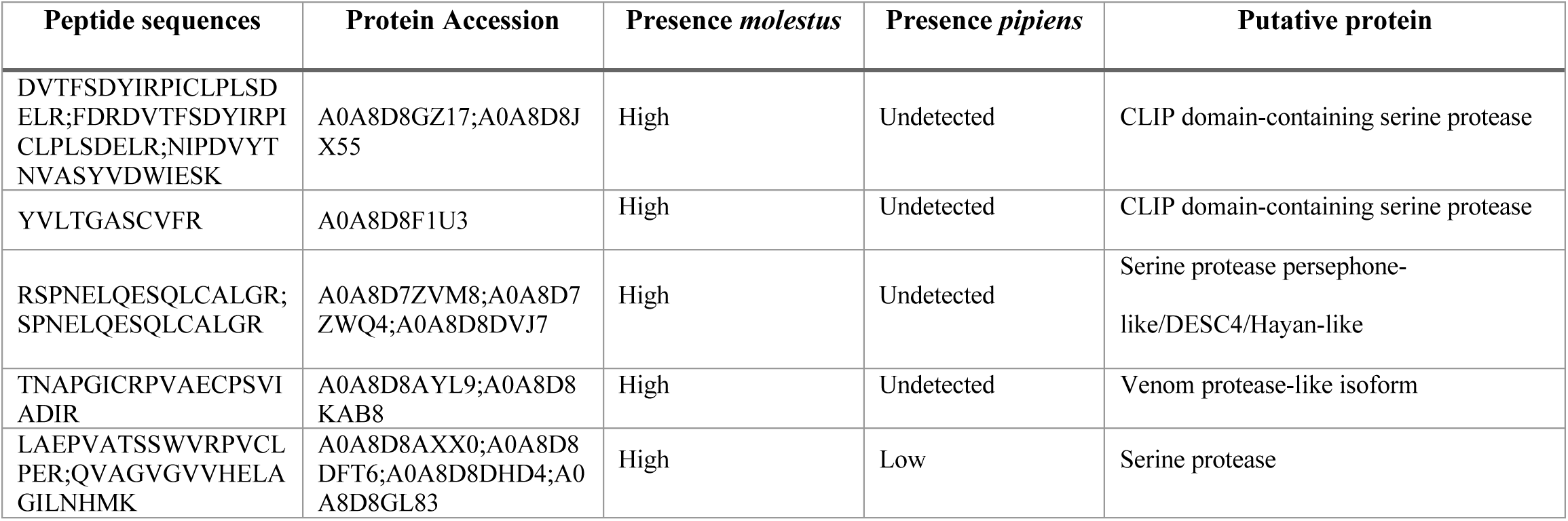
Top 5 most differentially present peptides found in *Cx.pip.molestus* versus *Cx.pip.pipiens* salivary gland extract in mass spec pilot analysis.

## Acknowledgements

The authors would like to thank Ingeborg van Middelkoop, Nicole Lambo, Dennis Akkermans, and Josianne Theuns – van Vliet for their assistance with the animal experiments; Wouter Doff for his help with the mass spectrometry; Pascal Miesen, Ezgi Taşköprü for their help with obtaining mosquito saliva; Pieter Rouweler and the insect rearing group from the Laboratory of Entomology at Wageningen University & Research for maintaining and providing the *Culex* mosquitoes.

